# Single-cell virology: On-chip, quantitative characterization of the dynamics of virus spread from one single cell to another

**DOI:** 10.1101/2024.09.25.615011

**Authors:** Wu Liu, Claus O. Wilke, Jamie J. Arnold, Mohamad S. Sotoudegan, Craig E. Cameron

## Abstract

Virus spread at the single-cell level is largely uncharacterized. We have designed and constructed a microfluidic device in which each nanowell contained a single, infected cell (donor) and a single, uninfected cell (recipient). Using a GFP-expressing poliovirus as our model, we observed both lytic and non-lytic spread. Donor cells supporting lytic spread established infection earlier than those supporting non-lytic spread. However, non-lytic spread established infections in recipient cells substantially faster than lytic spread and yielded higher rates of genome replication. While lytic spread was sensitive to the presence of capsid entry/uncoating inhibitors, non-lytic spread was not. Consistent with emerging models for non-lytic spread of enteroviruses using autophagy, reduction of LC3 levels in cells impaired non-lytic spread and elevated the fraction of virus in donor cells spreading lytically. The ability to distinguish lytic and non-lytic spread unambiguously will enable discovery of viral and host factors and host pathways used for non-lytic spread of enteroviruses and other viruses as well.

## Introduction

The use of the plaque assay in virology applied a strong selective pressure for the use of cell lines that cause substantial cytopathic effect and cell death in response to infection [1–2]. For non-enveloped viruses like the enteroviruses, this circumstance also led to the general conclusion that these viruses use a lytic mechanism for egress, with the inference that such a mechanism of spread might also occur in the infected host. The capacity of some cell lines to support persistent infection of enteroviruses provided the earliest hint that non-lytic mechanisms for egress must exist [3–6]. It is now abundantly clear that many non-enveloped viruses spread by non-lytic mechanisms [7–8]. Elaborating the details of these non-lytic pathways represents an exciting, ever-expanding area of virology.

Non-lytic spread by poliovirus is among the best described, although numerous gaps remain [9–14]. Infection induces macroautophagy (autophagy) by an ill-defined, unconventional mechanism based on the observation that the class-3 phosphatidylinositol-3 kinase (PI3KC3) that is essential for initiation of homeostatic autophagy is not required for PV-induced autophagy [15–16]. Infectious viral particles are thought to be incorporated into autophagosomes in a manner dependent on microtubule-associated protein 1A/1B-light chain 3 protein (LC3) [11–12] consistent with homeostatic mechanisms [15]. Autophagosomes containing viral particles bypass fusion with lysosomes because infection leads to degradation of the protein responsible for tethering autophagosomes to lysosomes [11,12,15,17]. Evasion of the lysosome leaves the plasma membrane as the target for fusion of autophagosomes, leading to secretion of vesicles containing infectious particles, also referred to as quasi-enveloped particles [18]. Uptake of these vesicles by recipient cells occurs; however, the mechanism is not clear [9]. One study has suggested that uptake requires the PV receptor [13].

Major confounders of studies of PV spread at the population level specifically, or any virus in general, include the between-cell heterogeneity in infection dynamics [19–22] that likely extrapolates to the dynamics of virus release and release of both free and quasi-enveloped virus into the same culture medium. The first and best effort to address these challenges visualized autophagy, infection, and several parameters linked to cell viability by time-lapse microscopy [23]. In this study, infection of recipient cells occurred prior to death of donor cells, consistent with non-lytic spread [23]. The assumption was that donor-recipient relationships could be inferred by proximity, which was likely the case but not absolute. Therefore, room exists for development of even more robust platforms to study virus spread.

Our laboratory developed a microfluidics platform for high-throughput investigation of PV infection dynamics in isolated single cells [19]. By adding a single cell-trapping structure to each nanowell, nanowell occupancy approached 90% [20]. Here, we report the design and construction of a device capable of trapping two different cells in each nanowell. With this tool, we demonstrate heterogeneity in the outcomes of primary infection from the perspective of the recipient cells: no spread, lytic spread, or non-lytic spread. Lytic spread was sensitive to an inhibitor targeting the PV capsid required, but non-lytic spread was not. Secondary infections established by non-lytic spread were more intense—that is, higher multiplicity of infection and genomes produced, than those established by lytic infection. Finally, non-lytic spread was PIK3C3-independent and LC3-dependent, consistent with current knowledge for PV. We conclude that the experimental paradigm presented will enable discovery of viral and host determinants of spread and enumeration of the pathway(s) taken by a viral particle from formation to release.

## RESULTS

### Observation and quantitation of modes of PV spread at single-cell resolution

By addition of two cell-trapping structures in each nanowell separated by a distance permitting each side of the device to be loaded independently, we were able to trap pairs of donor (infected) and recipient (uninfected) cells (**Figure 1** and **Figure S1**). On each device, 800 microwells were fabricated and ∼500 cell pairs could be isolated under optimized conditions.

**Figure 1.**
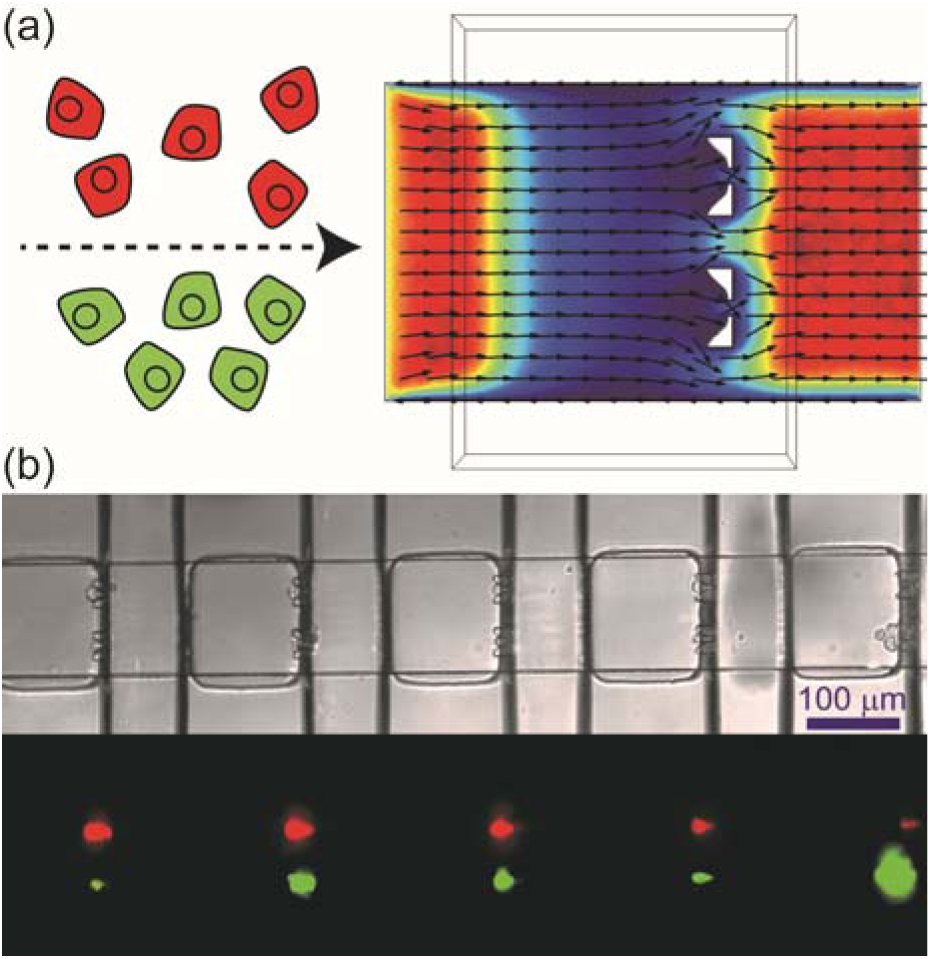
Cell pairing for unambiguous characterization of the spread of virus infection at the single-cell level. (**a**) Utilizing laminar flow in microchannels, a microfluidic device has been developed to capture two different cells in each nanowell. Vybrant DiI-labeled cells (red) and DiO-labeled cells (green) were infused into the two inlets of the channel at the same flow rate. Streamlines obtained by COMSOL simulation of the velocity field for the flow in the channel indicated that the two cell suspensions would not mix with each other. (**b**) Brightfield and fluorescence images show red cells captured on one side while green cells were captured on the other side of the same nanowell. Optimized condition: infusion rate was 100 nL/min; cell density was 8 × 10^5^ cells/mL.

In order to unambiguously visualize the process of spread, the GFP gene was fused in frame between the P1- and P2-coding regions of the PV genome as a reporter of polyprotein translation and infection, and the donor cells were labeled red using Vybrant DiI. We will refer to this reporter virus as PV-GFP. The cell pairs were imaged every 30 minutes from one to 24 hours post-infection (hpi) (**Figure 2**). In this work, PV-GFP at an MOI of 5 PFU/cell were used for adsorption to the donor cells, 72% ± 3% of which turned from red to yellow as a result of establishment of infection and viral replication. By 22 hpi, approximately half of the viral infections in donor cells failed to spread to the recipient cells (**Figure 2a** and **Movie S1**). For the other half, spread was marked by production of the GFP signal in the recipient cell. In addition to spread via the well-known lytic mode, (**Figure 2b** and **Movie S2**), we observed the onset of GFP expression in recipient cells as early as 11 hours prior to lysis of donor cells, providing direct observation of non-lytic spread by PV (**Figure 2c** and **Movie S3**).

**Figure 2.**
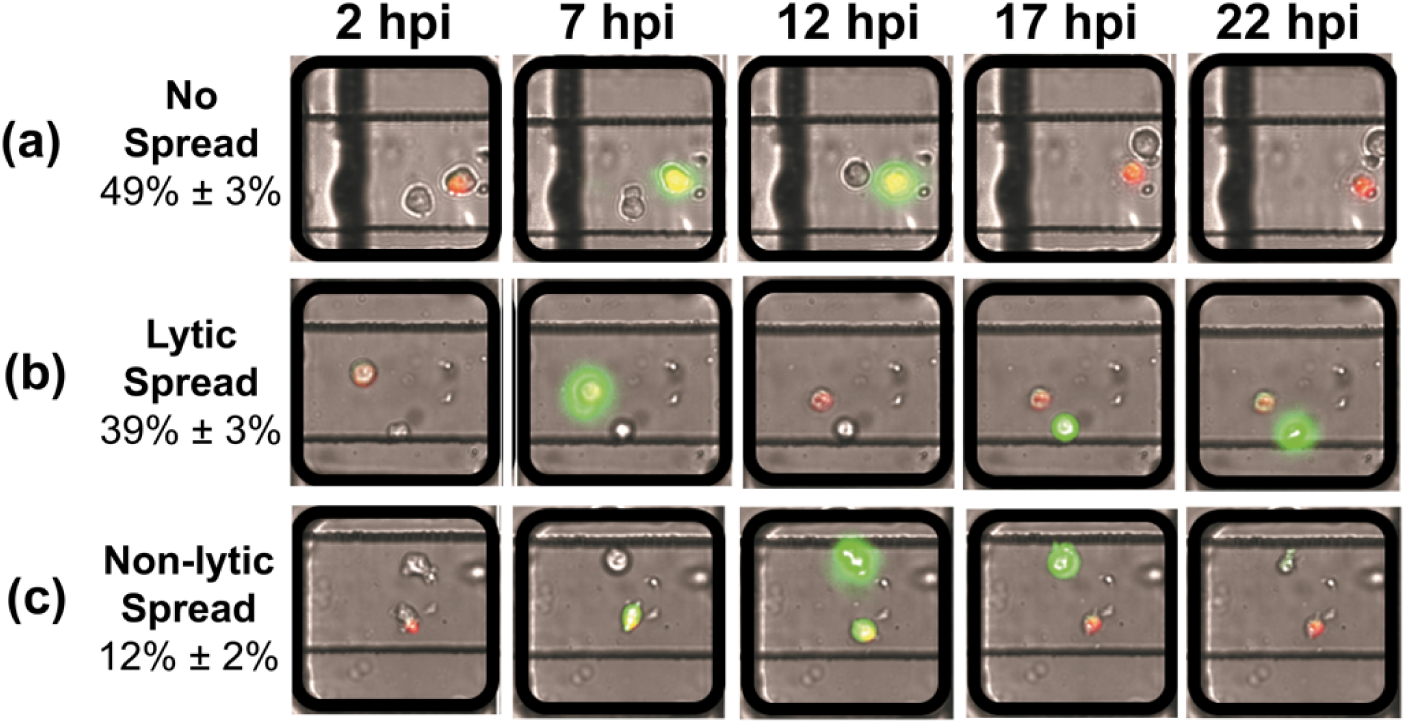
Different modes of PV spread observed by time-lapse imaging of cell pairs. Vybrant DiI (red) labeled cells were infected with PV-GFP (donor) and loaded simultaneously on the device with uninfected cells (recipient). The chip was imaged every 30 minutes over a period of 24 hours. Selected images are shown. As virus replicated in donor cells, green fluorescence was detected as yellow because of the red dye-labeled cell. As recipient cells became infected, these cells turned green. Lysis of infected donor cells caused the color to change from yellow to red as GFP leaked into the media. Three modes of spread were observed. (**a**) **No spread**: GFP signal increased in donor cells over time followed by lysis, indicated by the return of a red color to the donor cell, without any detectable GFP signal observed in recipient cells. (**b**) **Lytic spread**: GFP signal increased in donor cells over time followed by lysis, indicated by the return of a red color to the donor cell, followed later by observation of an ever-increasing GFP signal in recipient cells. (**c**) **Non-lytic spread**: GFP signal increased in donor cells over time; however, prior to lysis, indicated by the absence of red donor cells, GFP signal was observed in recipient cells. In most cases, lysis of the donor cell eventually occurred as donor cells turned red.

For quantitative analysis, we extracted the fluorescence intensity from each single cell. As documented previously, infection is described using four phenomenological parameters: maximum, slope, start point, and infection time. These parameters represent the yield of replicated RNA as reflected by GFP expression, rate of replication, the time in which the GFP signal increases to a detectable level, and the time it takes for GFP fluorescence to go from the first detectable signal to its maximum value, respectively [24]. By comparing the start point in the recipient cell to the lysis time of the donor cell, we were able to distinguish lytic and non-lytic spread of PV. We used a time difference (Δt) of 1 h as the cutoff, as it takes at least 1 h for GFP fluorescence to be detected after infection (Figure 3a). Based on this criterion, by 21 hpi, 12% ± 2% of the PV infection events achieved spread via a non-lytic mechanism while 39% ± 3% via a lytic mechanism.

**Figure 3.**
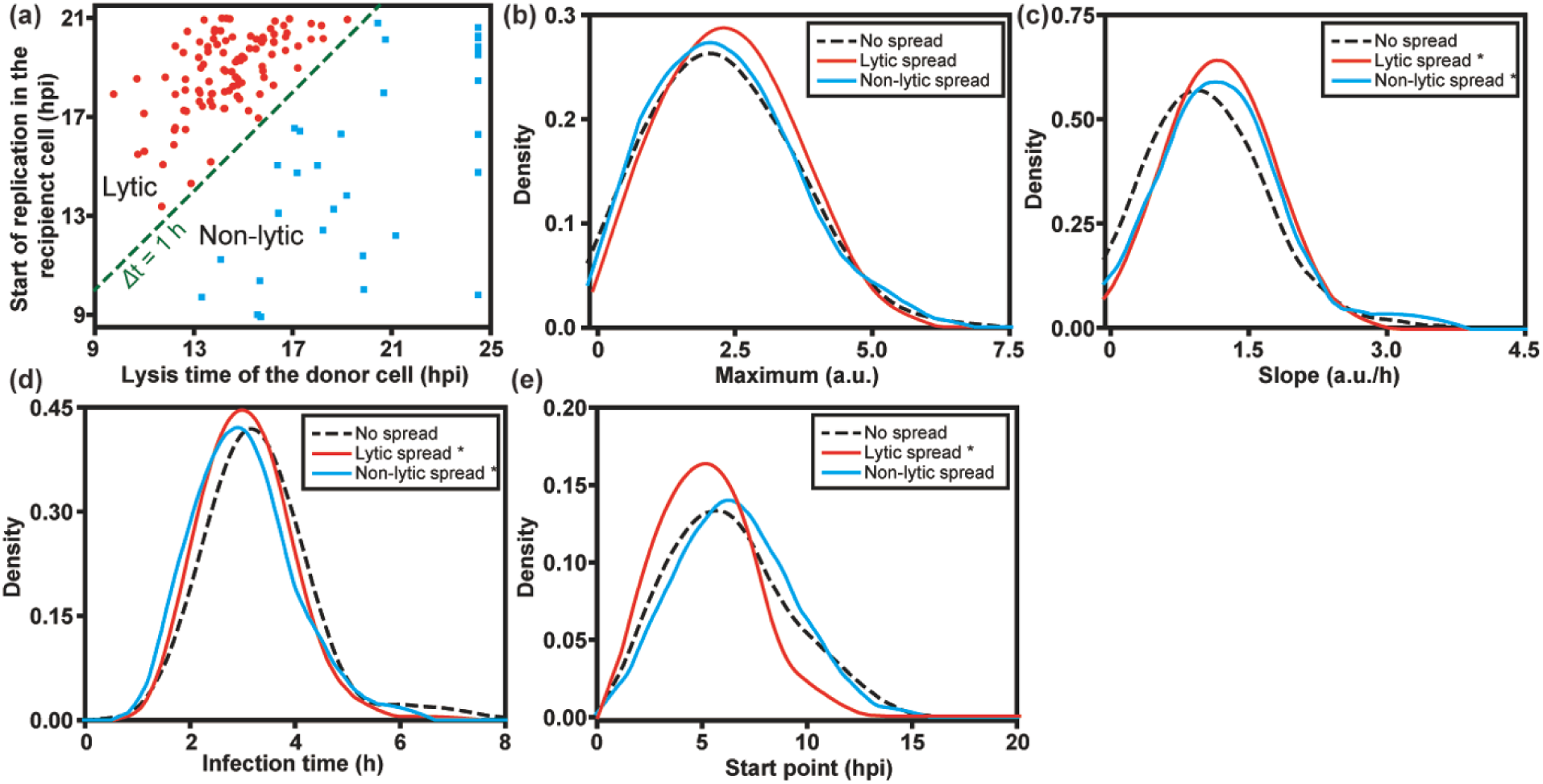
Quantitative analysis of PV infection dynamics in donor cells. (**a**) To distinguish between lytic and non-lytic spread, we represented each event in each well as the start time of replication in the recipient cell on the y-axis and the lysis time of the donor cell on the x-axis. A difference (y-x) greater than zero indicates lytic spread while a value less than zero indicates non-lytic spread. For donor cells that did not lyse by the end the experiment (24 hpi), 24.5 hpi was set as the lysis time here. To determine if infection dynamics in the donor cell influenced the mode of spread, distributions for experimental parameters measured in donor cells were plotted for each category of spread: no spread (**--**); lytic spread (--); and non-lytic spread (--). The following experimental parameters are shown: (**b**) maximum, (**c**) slope, (**d**) infection time, and (**e**) start point. *: p < 0.05 compared to no spread group based on t-test (**Table S1**).

We also compared the replication kinetic parameters in donor cells that failed to spread with donor cells that gave rise to lytic and non-lytic spread. As shown in **Figure 3b**, no significant differences in maximum were found among the groups. However, the speed of replication is faster in donor cells with either lytic or non-lytic spread, than in those without spread (**Figures 3c** and **3d**). In addition, infections leading to lytic spread also started earlier, which was not observed for non-lytic spread (compare **Figures 3c** and **3d**). We conclude that fast replication speed in host cells is a favorable factor for spread, perhaps to outpace intrinsic defenses of the cell.

### Hallmarks of non-lytic spread: fast, high multiplicity, reduced sensitivity to capsid-binding inhibitors

To understand if there are more differences between the lytic and non-lytic modes of spread beyond the time point in which virions are released, we compared the changes in the kinetic parameters from the donor cell to the recipient cell in pairs occupying the same nanowell. We refer to the interval between start points in the donor and recipient cells as the “cycle time” for spread. It typically required 10–18 h for PV infection to spread lytically from one cell to another (**Figure 4a**). For non-lytic spread, an average of ∼10 h with a wide variance was observed, consistent with non-lytic spread being the fastest mode of PV transmission from one cell to another. For both non-lytic and lytic spread, quantitative measures of outcomes of infection in recipient cells exceeded corresponding measures in donor cells for yield (maximum) and replication speed (slope) (**Figures 4b** and **4c**). In several instances, the increase in yield and replication speed observed for non-lytic spread exceeded that observed for lytic spread by as much as 5- to 10-fold (**Figures 4b** and **4c**). Non-lytic spread clearly has the potential to be far more efficient than lytic spread. Interestingly, the infection time was independent of the cell (donor or recipient) or the mode of spread (non-lytic or lytic) (**Figure 4d**).

**Figure 4.**
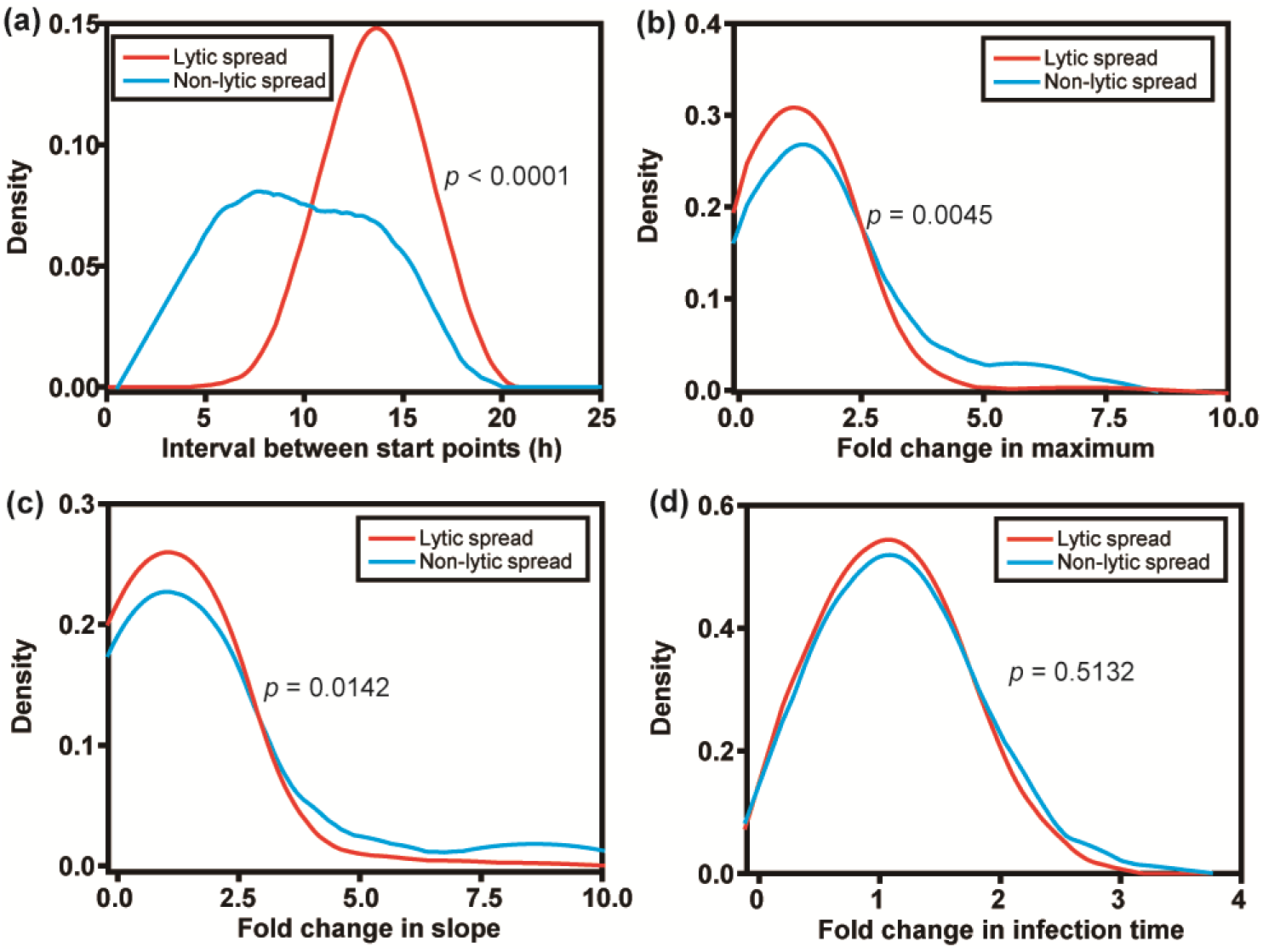
Quantitative analysis of PV infection dynamics in recipient cells: comparison of lytic and non-lytic spread. (**a**) Distributions of the difference (recipient-donor) for start point. Distributions of the fold change (parameter in recipient cell/parameter in donor cell) for (**b**) maximum, (**c**) slope, and (**d**) infection time.. In each case, lytic infections and non-lytic infections were treated independently. Non-lytic spread has the capacity to reach values of maximum higher than those observed for lytic spread in the same time period. Non-lytic spread appears to establish infection in recipient cells much faster than lytic spread.

Non-lytic carriers of PV are thought to have a single bilayer that would limit exposure of the internally localized capsids to substances present in the extracellular milieu [13]. Pocapavir is a small molecule that binds to the capsid of several enteroviruses and inhibits entry and/or uncoating [25]. Treatment of donor cells with pocapavir caused a substantial reduction in lytic spread at 1 µM, with near-complete elimination of lytic spread at 14 µM (**Figure 5a**). Under these conditions, non-lytic spread was minimally impacted (**Figure 5a**). Interestingly, infections that were able to withstand the sub-lethal dose of pocapavir elicited outcomes suggesting increased fitness of the infecting population relative to the virus that did not spread or spread by a non-lytic route (**Figures 5b** – **5e**). For statistical significance of the ovserved differences, p-values are presented in **Table S2**.

**Figure 5.**
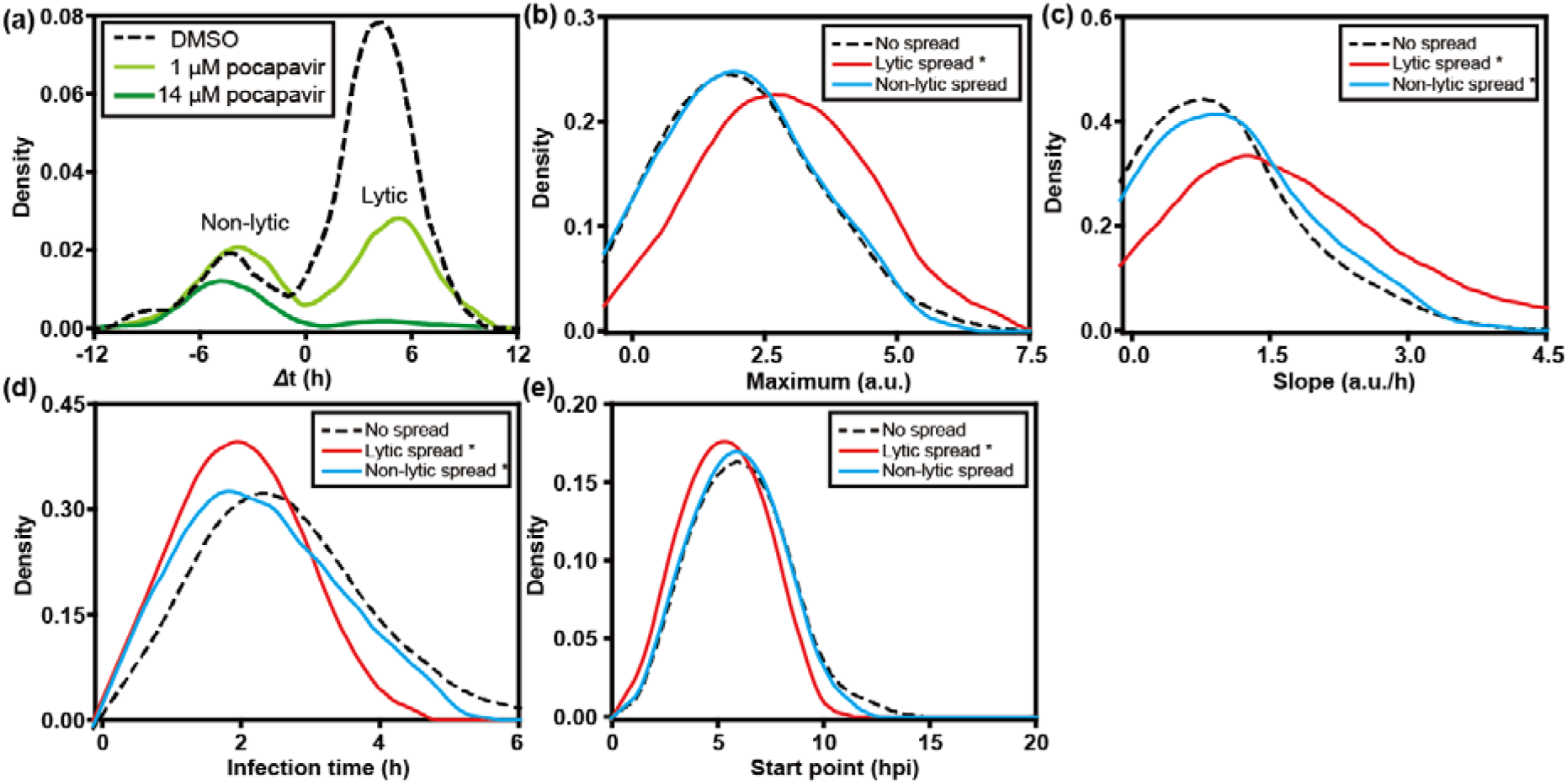
Lytic spread is sensitive to the presence of an entry/uncoating inhibitor; non-lytic spread is not. Pocapavir is known to bind to the PV capsid, which interferes with entry or uncoating. We have evaluated the impact of the presence of pocapavir on establishment of PV infection in recipient cells after lytic or non-lytic spread. (**a**) Infected, donor cells were added to uninfected recipient cells in the absence or presence of the indicate concentration of pocapvir. The impact of the drug on spread was evaluated. The area under each curve is proportional to the number of events observed. Lytic spread was sensitive to the presence of pocapavir. In order to determine how pocapavir (1 µM) impacted each parameter, we evaluated distributions for the following parameters: (**b**) maximum, (**c**) slope, (**d**) infection time, and (**e**) start point in the donor cells stratified by spread phenotype: no spread (**--**); lytic spread (--); and non-lytic spread (--). Higher values of maximum and slope, and earlier start time of infection in donor cells that lead to lytic spread suggests that viruses that are resistant to the capsid inhibitor may have higher viral fitness.

### Role of autophagy in non-lytic spread of PV

It is now widely accepted that non-lytic spread by PV specifically, and perhaps enteroviruses broadly, exploits a specialized, virus-induced form of secretory autophagy [11]. Induction of normal, cellular autophagy requires Vps34, a class III PI3 kinase that produces PI3P [26]. PV-induced autophagy does not [16]. Importantly, non-lytic spread of PV observed here was also independent of Vps34 as non-lytic spread was not sensitive to the presence of SAR405, an inhibitor of Vps34 (**Figure 6**). In contrast, LC3B is required for PV-induced autophagy and knockdown of LC3B expression by using siRNA in the donor cells (**Figure S2**) reduced substantially the frequency of non-lytic spread (**Figure 6**). Failure of virus to be released non-lytically led to an increase in the amount of virus that used a lytic mode of spread (**Figure 6**).

**Figure 6.**
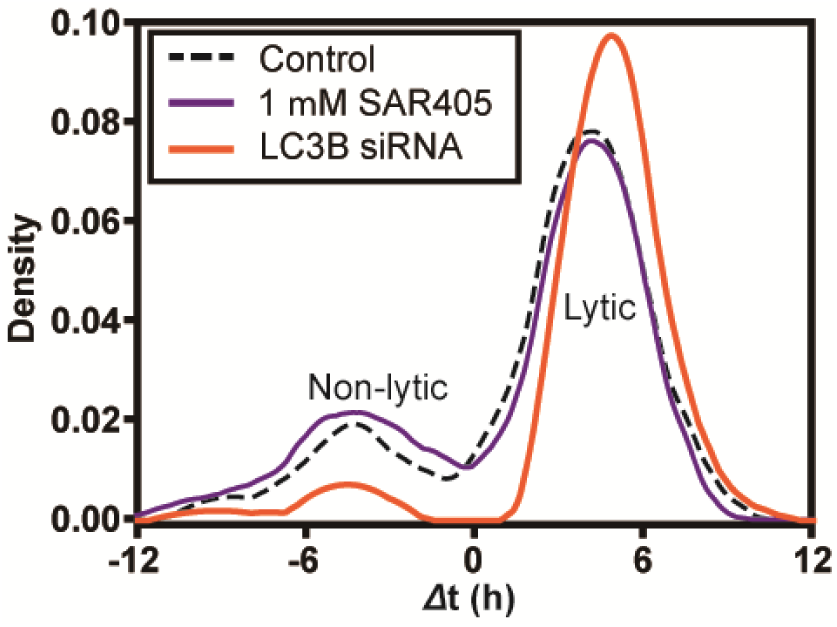
Non-lytic spread between single cells uses a non-traditional mechanism of autophagy as observed in bulk culture. Non-lytic spread of enteroviruses uses secretory autophagy, which is dependent on LC3 but independent of Vps34 (target of SAR405). To assess the relationship of non-lytic spread observed between single cells to that observed at the population level in cell culture, we evaluated the requirement for Vps34 by monitoring spread in the presence of SAR405 or from donor cells in which LC3B had been depleted (LC3B siRNA). SAR405 exhibited no impact on spread. Loss of LC3B interfered selectively with non-lytic spread.

In conclusion, by experimentally discriminating the two modes of spread, we have quantitatively characterized the dynamics of the lytic and nonlytic modes of PV infection at the single-cell level (**Figure 7**). Although only giving rise to a small proportion of spread events in the experiments performed here, the non-lytic mode exhibits stronger enhancement in viral replication dynamic parameters, suggesting its potential role in efficient and rapid spread of viral infection. The presented strategy to investigate the dissemination of viral infection, which is inaccessible through conventional experimental approaches, would promote the elucidation of the unclear mechanism of non-lytic spread and the discovery of targets to block non-lytic spread of enteroviruses.

**Figure 7.**
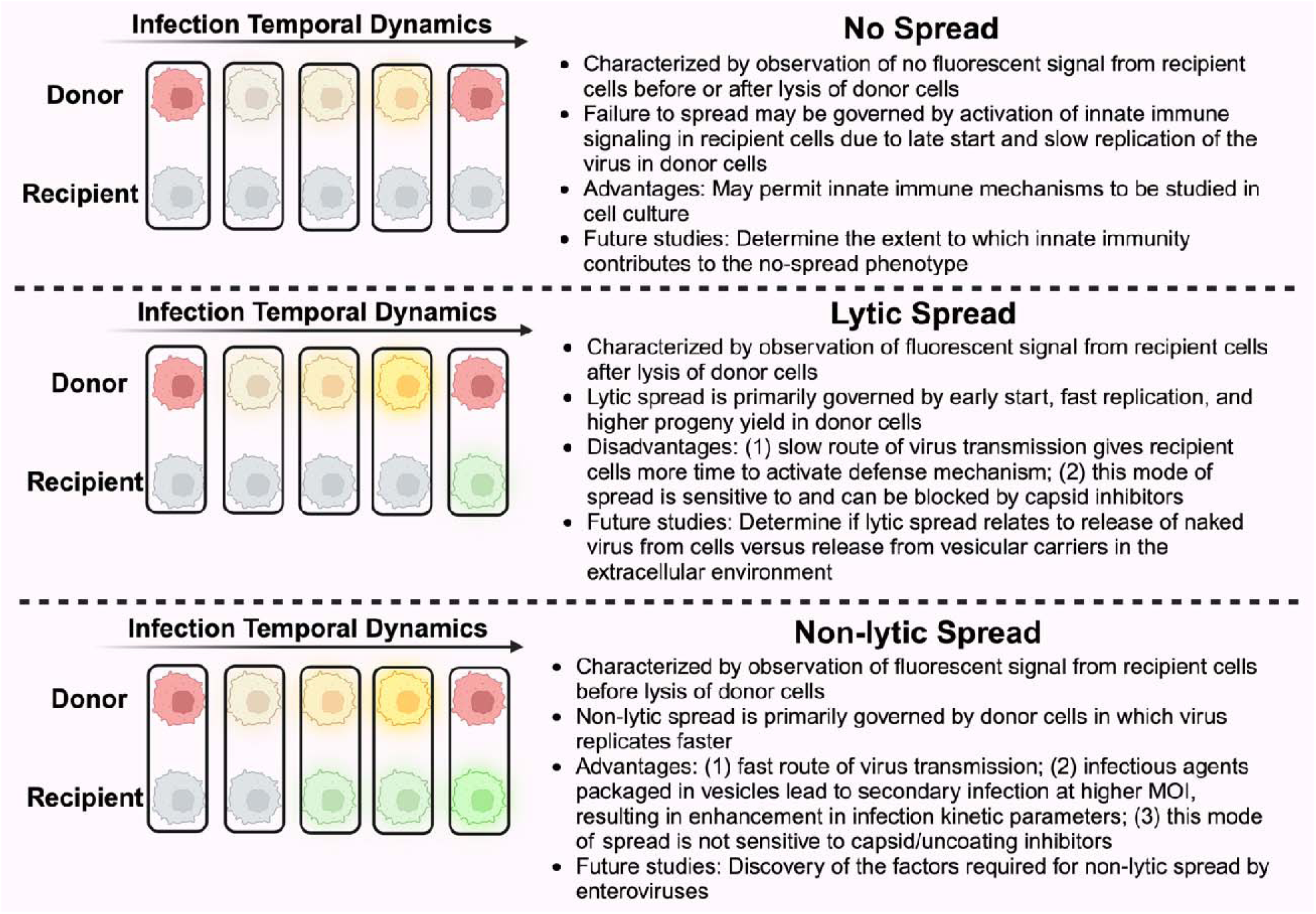
Summary of virus-spread phenotypes observed in this study and corresponding implications. Primary infection is charazterized by observation of yellow fluorescence signal in donor cells (stained red at *t* = 0). Lysis is marked by loss of yellow signal and return of red signal. Observation of a green signal from recipient cells before or after lysis of donor cells distinguishes between lytic and non-lytic spread. The absence of green fluorescence in recipient cells indicates failure of infection to spread. Governing factors, advantages or disadvantages, and future studies associated with each mode of spread are described.

## DISCUSSION

How a virus or population of viruses moves from one cell to another is a fundamental principle of infection biology. There was a time when virologists considered virus spread as a mutually exclusive, binary process: either lytic or non-lytic. In recent years, however, it has become increasingly clear that lytic spread may reflect a mechanism used in certain cell-based models as opposed to being representative of the primary mechanism of spread in humans or animals. For poliovirus (PV) and other enteroviruses, the pioneering work of the Kirkegaard laboratory motivated virologists to begin to think about alternative, non-lytic mechanisms of virus spread for viruses thought to spread by a lytic mechanism [9–10,23]. There is now substantial evidence supporting the hijacking of the cellular secretory autophagy pathway by enteroviruses to create vesicular carriers of enterovirus virions [8,10–12,14,16–17,23]. Because a lytic or a non-lytic mechanism occurs in any given cell in culture, robust, rigorous approaches to study viral spead in culture has been at best challenging [23], if not impossible.

Our laboratory has had a longstanding interest in viral infection dynamics on the single-cell level [19–20]. The primary motivation for this study was to ask whether or not virus transmission from one single cell to another could be studied using technology similar to that used for our single-cell-virology experiments [19–20]. Such an advance would make it possible to begin to dissect the similarities and differences between lytic and non-lytic spread. Further, if non-lytic spread could be observed, then it would be possible to dissect the viral and host factors contributing to this form of spread. By adding two traps to each well of the microfluidics device used for our traditional single-cell-virology experiments (**Figure 1** and **Figure S1**), we were able to observe three modes of spread: no spread, lytic spread, and non-lytic spread (**Figure 2**). Each mode of spread exhibited a unique kinetic signature when evaluating viral spread dynamics from the infected, donor cell to the uninfected, recipient cell (**Figure 3**).

The first major finding of this study is unambiguous evidence for the ability of individual HeLa cells from a single population of cells to support different modes of spread (**Figure 2**). Demonstration of such heterogeneity highlights the complication of studying spread in bulk culture. However, a lot of fun will be had deciphering the viral and/or host contributions to this heterogeneity. The biggest surprise, however, was the observation that half of the primary, lytic infections failed to establish a secondary infection (**Figure 2a**). Upon inspection of the parameters used to assess infection dynamics at the single-cell level (**Figure 3**), the most distinguishing feature of this mode of spread (no spread) was that infection in the primary cell was established “late” relative to primary infections that lead to other modes of spread (**Figure 3e**). Our current hypothesis is that establishment of infection “late” may actually permit activation of intrinsic defenses in the infected cell and release of interferon or other signaling molecule that induces an effective antiviral response in the uninfected, recipient cells. If this is the case, then this system could prove useful in dissecting host responses to viral infection.

When we compare lytic spread to non-lytic spread, the data suggest that lytic spread occured more often than non-lytic spread (**Figure 2** and **Figure 3a**). In our system, the two modes of spread are distinguished by the time green fluorescence appears in the uninfected, recipient cell relative to the time in which green fluorescence diminishes in the infected, donor cell (**Figures 2b** and **2c**). Therefore, a major limitation of our study is the use of GFP, which has a very long and heterogeneous activation time [27]. It is possible that virus could spread using a non-lytic mechanism but that development of the fluorophore in the recipient cell fortuitously occured after the donor cell lysed. Going forward, it will be important to try a fluorophore that forms immediately upon expression to determine the extent to which the kinetics of flurophore activation impacts interpretation of spread dynamics.

When PV moves by lytic spread, the cell becomes permeable to GFP but remains intact (**Movie S2**). Does lytic spread of PV use these permeable regions of the plasma membrane? Is it possible that mechanisms that permeabilize the plasma membrane also permeabilize the vesiclular carriers and thereby cause release of virus? Is it possible that lytic spread reflects slow disruption of vesicular carriers present in the media? There is clearly a relationship between lytic and non-lytic spread as interference with non-lytic spread leads to an increase in virus moving by a lytic mechanism (**Figure 6**). More work will be required to delineate the rules of lytic spread and non-lytic spread and the relationships between the two modes of spread. The approach reported here should be useful in a careful dissection of these modes of spread.

This study revealed a major difference between lytic and non-lytic spread that had not been considered when the study was originally contemplated. Lytic spread of PV in cell culture is well known to be inhibited by molecules that bind the capsid, impairing entry and/or uncoating. This phenomenon was observed at the single-cell level as well as using pocapavir (**Figure 5a**). Interestingly, the naked virions that survived challenge by pocapavir exhibited enhanced fitness relative to typical member of the virus population (**Figures 5b** - **e**). Importantly, non-lytic spread was not sensitive to pocapavir (**Figure 5a**). How vesicular carriers of PV enter the cell is unclear, but there is evidence that the PV receptor is required for such carriers to establish infection [13]. The inability for capsid inhibitors to interfere with PV virions in vesicular carriers has interesting implications. If a small molecule cannot access the inside of the vesicular carriers, then it is likely that these carriers will protect virions from immune cells, antibodies, and other strategies that have evolved to address pathogens in circulation. A final benefit of the vesicular carriers revealed by this study is that the intensity of the infection established by these carriers is greater than naked virions (**Figure 4**), consistent with vesicular carriers promoting a higher multiplicity of infection and the potential for cooperative interactions between co-infecting virions [28].

The final question we asked was whether or not the non-lytic spread we observe at the single-cell level is related to the autophagy-dependent process observed in bulk culture [11]. Non-lytic spread was dependent on LC3B (**Figure 6**) as previously observed [23] and used a Vps34-independent mechanism (**Figure 6**) also as previously observed [16]. These initial results are consistent with the between-cell spread observed in nanowells representing that observed in bulk cell culture.

This study has demonstrated the utility of the cell-pairing device for the study of viral spread. However, there is great potential for the cell-pairing concept to be applied to other aspects of infection biology, particularly, intercellular signaling and immune cell responses to infected cells. For example, exosomes or exosome-like vesicles can be released from infected cells and initiate proviral changes to uninfected, bystander cells [29]. With fluorescent reporters for activation or suppression of the the appropriate intracellular pathway(s), the kinetics and heterogeneity of the response of a cell to released exosomes can be investigated. There are numerous cell types of the immune system that can interact with the infected cell [30]. Macrophage phagocytose infected cells. CD8-positive T cells recognize viral peptides presented on the surface of the infected cell, triggering responses that lead to death of the infected cell. The cell-pairing device should prove useful for the study of these and other immune-cell-driven interactions with infected cells. The primary barrier to completing these studies at present is the absence of optical signals to use to monitor these interactions, so some biological engineering will be required.

## MATERIALS AND METHODS

### Cells, viruses and reagents

HeLa S3 cells were obtained from American Type Culture Collection (ATCC) and maintained in DMEM/F12 (1:1) (Life Technologies), supplemented with 10% fetal bovine serum (FBS, Atlanta Biologicals) and 100 IU/mL penicillin-streptomycin (Corning), in a humidified atmosphere of 95% air and 5% CO_2_ at 37 °C.

eGFP-tagged poliovirus was prepared as previously described [19–20], despite that the eGFP sequence was modified to improve virus specific infectivity. Briefly, PV-eGFP plasmid was linearized with ApaI and purified with QIAEX II Gel Extraction Kit (Qiagen, Netherlands). Viral RNA was then transcribed at 37°C for 5.5 hours in a 20-μL reaction medium containing 350 mM HEPES pH 7.5, 32 mM magnesium acetate, 40 mM dithiothreitol (DTT), 2 mM spermidine, 28 mM nucleoside triphosphates (NTPs), 0.025 µg/µL linearized DNA, and 0.025 µg/µL T7 RNA polymerase. Magnesium pyrophosphate in the mixture was removed by centrifugation. Qiagen RNeasy kit was used to purify viral RNA. 1 µg of viral RNA were utilized to transfect HeLa cells cultured at 37°C using TransMessenger reagent (Qiagen) following the manufacturers protocol. PV-eGFP was harvested by three repeated freeze-thaw cycles, and suspended in 0.5% nonidet P-40 (NP-40). For purification, 1 volume of 20% PEG-8000/1 M NaCl solution was added. The mixture was incubated overnight at 4°C and the supernatant was removed by centrifugation at 8000 × g for 10 minutes at 4°C. The pellet (virus) was then resuspended in PBS and filtered with Centricon® Plus-70 (EMD Millipore, USA). Virus titer was determined by plaque assay.

Pocapavir and SAR405 were purchased from MedChemExpress (Monmouth Junction, NJ).

### RNA Interference

A pool of LC3B targeting or non-targeting control siRNAs (Dharmacon) was transfected into HeLa S3 cells in 24-well plates according to manufacturer’s protocol. One day prior to transfection, 2.5 × 10^4^ cells were seeded in each well and incubated overnight. 12.5 pmol of siRNAs and 2 μL of DharmaFECT reagent were diluted in 50 μL of serum-free DMEM-F12 respectively, and incubated at room temperature for 5 min. Diluted siRNAs and DharmaFECT reagent were then mixed and incubated at room temperature for 20 min. 400 μL of complete medium was then added to the transfection complexes and added to the well.

### Western Blotting

At 72 hours post-transfection (hpt), cells were collected and lysed using RIPA buffer (10 mM Tris-HCl pH 8.0, 1 mM EDTA, 1 mM EGTA, 1 % Triton X-100, 0.1 % sodium deoxycholate, 0.1 % SDS, 140 mM NaCl, and 1 mM PMSF). Lysate protein concentration was determined by Bradford assay. 60 µg was used for SDS-PAGE and transferred onto nitrocellulose membrane. Membrane was blocked in 5% milk in TBST (20 mM Tris-HCl pH7.6, 137 mM NaCl, and 0.1 % Tween-20) and incubated with rabbit anti-LC3B primary antibody (Abcam, ab192890) and mouse anti-beta actin primary antibody (Abcam, ab6276), followed by incubation with goat anti-rabbit secondary antibody conjugated with alkaline phosphatase (KPL secondary Antibodies, Seracare) and the fluorescence signals were visualized using ECF substrate (GE Healthcare, RPN2785) in spectrometer (G:Box, Syngene).

### Microfluidic devices

The microfluidic devices designed to pair two different cells (Figure S1) were fabricated from polydimethylsiloxane (PDMS, GE RTV615) as described previously [19–20]. Briefly, SU-8 layers on silicon wafers were fabricated with desired thicknesses by soft lithography. Valve layer was generated by curing premixed PDMS prepolymer and curing agent (ratio 5:1). For channel layer, premixed PDMS prepolymer and curing agent (ratio 20/1) were spin-coated on the mold for 1 minute at 1500 rpm and incubated at 65 °C for 20 min. The valve layer was then released from its mold and aligned with the channel layer. The assembly was incubated overnight at 65 °C. The assembly was released from the mold and further bonded with alignment to the microwells layer, which was generated by curing premixed PDMS prepolymer and curing agent (ratio 20/1), by baking at 65 °C overnight. The obtained device was further bonded to a glass slide.

### On-chip experiments

HeLa S3 cells were labeled by Vybrant DiD, mixed with PV-GFP at the MOI of 5 PFU/cell, and shaken at 140 rpm for 30 min. The cells washed with PBS for three times, and resuspended to a density of 8 × 10^5^ cells/mL in normal culture medium. This infected-cells suspension and a suspension of un-infected cells were simultaneously infused to the two inlets of a microfluidic channel, both at 200 nL/min for 5 min. A pressure of 30 psi was then applied to the valves to isolate the microwells.

For microscopic imaging, the microfluidic device was placed in the chamber of a WSKM GM2000 incubation system (Tokai, Japan), which was adapted to a Nikon Eclipse Ti inverted microscope (Nikon, Japan). Automatic bright-field and fluorescence imaging were performed every 30 minutes from 3 hpi to 24 hpi with a ProScan II motorized flat top stage (Prior Scientific, USA), a CFI60 Plan Apochromat Lambda 10× objective, and a Hamamatsu C11440 camera.

### Data processing

Fluorescence intensity of the single cells and background intensity of the microwells were extracted with a customized MATLAB script. Relative intensity was calculated as (*Cell intensity* - *Background*)/*Background*. The relative intensity over was thus obtained to describe the viral replication dynamics in the single cells. As documented in our previous papers, parameters *maximum*, *slope*, *infection time* and *start point* were derived [19–20].

## SUPPLEMENTARY MATERIALS

Design of the microfluidic device; distributions and between-group t-test of the parameters indicated; videos showing different modes of PV spread.

## ACKNOWLEDGEMENTS

This study was initiated as our laboratory was preparing to move to Chapel Hill, NC. Then, we had a pandemic. There were several members of the Single-Cell-Virology team who contributed to this study in ways that could have been acknowledged by authorship. However, the diaspora of our colleagies has made it difficult to find everyone. We thank Chia-Heng Hsiung, Andrew Woodman, Mehmet U. Caglar, and Cory D. DuBois for their contributions to enabling this study. W.L. thanks Dr. Siyang Zheng for sharing the equipment to fabricate the microfluidic devices and Dr. Zhangming Mao for helpful discussions.

## AUTHOR CONTRIBUTIONS

Conceptualization, C.E.C., M.S.S, J.J.A., and W.L.; Methodology, C.E.C., M.S.S., and W.L.; Software, C.O.W., and W.L.; Validation, C.E.C., M.S.S., C.O.W., and W.L.; Formal Analysis, C.O.W.; Investigation, C.E.C., M.S.S., and W.L.; Resources, C.E.C., and C.O.W.; Data Curation, C.O.W., and W.L.; Writing – Original Draft Preparation, C.E.C., and W.L.; Writing – Review & Editing, C.E.C., M.S.S., and J.J.A.; Visualization, M.S.S., and W.L.; Supervision, C.E.C., and M.S.S.; Project Administration, C.E.C.; Funding Acquisition, C.E.C., and J.J.A.

## REFERENCES

1. Dulbecco, R.; Vogt, M. Some problems of animal virology as studied by the plaque technique. Cold Spring Harb Symp Quant Biol. 1953, 18, 273–9.

2. 2. Cooper, P.D., The Plaque Assay of Animal Viruses. Advances in Virus Research, Smith, K.M., Lauffer, M.A., Eds.; Academic Press 1962, Volume 8, 319–378.

3. Nekoua, M.P., Alidjinou, E.K. & Hober, D. Persistent coxsackievirus B infection and pathogenesis of type 1 diabetes mellitus. Nat Rev Endocrinol 2022, 18, 503–516.

4. Kandolf, R., Klingel, K., Zell, R., Selinka, H. C., Raab, U., Schneider-Brachert, W., and Bültmann, B. Molecular pathogenesis of enterovirus-induced myocarditis: virus persistence and chronic inflammation. Intervirology 1993, 35(1-4), 140.

5. Lloyd, R. E., & Bovee, M. Persistent infection of human erythroblastoid cells by poliovirus. Virology 1993, 194(1), 200–209.

6. Pelletier, I., Duncan, G., & Colbère-Garapin, F. One amino acid change on the capsid surface of poliovirus sabin 1 allows the establishment of persistent infections in HEp-2c cell cultures. Virology 1998, 241(1), 1–13.

7. Feng, Z.; Hensley, L.; McKnight, K. L.; Hu, F.; Madden, V.; Ping, L.; Jeong, S. H.; Walker, C.; Lanford, R. E.; Lemon, S. M. A pathogenic picornavirus acquires an envelope by hijacking cellular membranes. Nature 2013, 496, 367–371.

8. Robinson, S. M.; Tsueng, G.; Sin, J.; Mangale, V.; Rahawi, S.; McIntyre, L. L.; Williams, W.; Kha, N.; Cruz, C.; Hancock, B. M.; Nguyen, D. P.; Sayen, M. R.; Hilton, B. J.; Doran, K. S.; Segall, A. M.; Wolkowicz, R.; Cornell, C. T.; Whitton, J. L.; Gottlieb, R. A.; Feuer, R. Coxsackievirus B exits the host cell in shed microvesicles displaying autophagosomal markers. PLoS Pathog. 2014, 10, e1004045.

9. Bird, S. W., & Kirkegaard, K. Escape of non-enveloped virus from intact cells. Virology 2015, 479, 444–449.

10. Bird, S. W., & Kirkegaard, K. Nonlytic spread of naked viruses. Autophagy 2015, 11(2), 430–431.

11. Jassey, A., & Jackson, W. T. Viruses and autophagy: bend, but don’t break. Nature Reviews Microbiology 2024, 22(5), 309–321.

12. Jackson, W. T. Viruses and the autophagy pathway. Virology 2015, 479, 450–456.

13. Chen, Y. H.; Du W; Hagemeijer, M. C.; Takvorian, P. M.; Pau, C.; Cali, A.; Brantner, C. A.; Stempinski, E. S.; Connelly, P. S.; Ma, H. C.; Jiang, P.; Wimmer, E.; Altan-Bonnet, G.; Altan-Bonnet, N. Phosphatidylserine vesicles enable efficient en bloc transmission of enteroviruses. Cell 2015, 160, 619–630.

14. Mutsafi, Y., & Altan-Bonnet, N. Enterovirus transmission by secretory autophagy. Viruses 2018, 10(3), 139.

15. Melia, T. J., Lystad, A. H., Simonsen, A. Autophagosome biogenesis: From membrane growth to closure. J. Cell Biol. 2020; 219(6):e202002085.

16. Corona Velazquez, A., Corona, A. K., Klein, K. A., & Jackson, W. T. Poliovirus induces autophagic signaling independent of the ULK1 complex. Autophagy 2018, 14(7), 1201–1213.

17. Corona, A. K., Saulsbery, H. M., Velazquez, A. F. C., & Jackson, W. T. Enteroviruses remodel autophagic trafficking through regulation of host SNARE proteins to promote virus replication and cell exit. Cell reports 2018, 22(12), 3304–3314.

18. Das, A., Rivera-Serrano, E. E., Yin, X., Walker, C. M., Feng, Z., and Lemon, S. M. Cell entry and release of quasi-enveloped human hepatitis viruses. Nature Reviews Microbiology 2023, 21(9), 573–589.

19. Guo, F.; Li, S.; Caglar, M. U.; Mao, Z.; Liu, W.; Woodman, A.; Arnold, J. J.; Wilke, C. O.; Huang, T. J.; Cameron, C. E. Single-cell virology: on-chip investigation of viral infection dynamics. Cell Rep. 2017, 21, 1692–1704.

20. Liu, W., Caglar, M. U.; Mao, Z.; Woodman, A.; Arnold, J. J.; Wilke, C. O.; Cameron, C. E. More than efficacy revealed by single-cell analysis of antiviral therapeutics. Sci. Adv. 2019, 5, eaax4761.

21. Dolan, P. T., Whitfield, Z. J., & Andino, R. Mapping the evolutionary potential of RNA viruses. Cell host & microbe 2018, 23(4), 435–446.

22. Dolan, P. T., Whitfield, Z. J., & Andino, R. Mechanisms and concepts in RNA virus population dynamics and evolution. Annual Review of Virology 2018, 5(1), 69–92.

23. Bird, S. W.; Maynard, N. D.; Covert, M. W.; Kirkegaard, K. Nonlytic viral spread enhanced by autophagy components. Proc. Natl. Acad. Sci. 2014, 111, 13081–13086.

24. Caglar, M. U.; Teufel, A. I.; Wilke, C. O. Sicegar: R package for sigmoidal and double-sigmoidal curve fitting. PeerJ, 2018, 6, e4251.

25. Oberste, M. S.; Moore, D.; Anderson, B.; Pallansch, M. A.; Pevear, D. C.; Collett, M. S. In vitro antiviral activity of V-073 against polioviruses. Antimicrob. Agents. Chemother. 2009, 53, 4501–4503.

26. Ronan, B.; Flamand, O.; Vescovi, L.; Dureuil, C.; Durand, L.; Fassy, F.; Bachelot, M. F.; Lamberton, A.; Mathieu, M.; Bertrand, T.; Marquette, J. P.; El-Ahmad, Y.; Filoche-Romme, B.; Schio, L.; Garcia-Echeverria, C.; Goulaouic, H.; Pasquier, B. A highly potent and selective Vps34 inhibitor alters vesicle trafficking and autophagy. Nat. Chem. Biol. 2014, 10, 1013–1019.

27. Balleza, E., Kim, J. M., & Cluzel, P. Systematic characterization of maturation time of fluorescent proteins in living cells. Nature methods 2018, 15(1), 47–51.

28. Sanjuán, R. The social life of viruses. Annual review of virology 2021, 8(1), 183–199.

29. Bello-Morales, R., Ripa, I., & López-Guerrero, J. A. Extracellular vesicles in viral spread and antiviral response. Viruses 2020, 12(6), 623.

30. Rouse, B. T., & Sehrawat, S. Immunity and immunopathology to viruses: what decides the outcome?. Nature Reviews Immunology 2010, 10(7), 514–526.

